# Flexible and transparent silver nanowire structures for multifunctional electrical and optical biointerfacing

**DOI:** 10.1101/2020.10.10.334755

**Authors:** Zhiyuan Chen, Nicolas Boyajian, Zexu Lin, Rose T. Yin, Sofian N. Obaid, Jinbi Tian, Jaclyn A. Brennan, Sheena W. Chen, Alana N. Miniovich, Leqi Lin, Yarong Qi, Xitong Liu, Igor R. Efimov, Luyao Lu

## Abstract

Transparent microelectrodes have recently emerged as a promising approach to combine electrophysiology with optophysiology for multifunctional biointerfacing. High-performance flexible platforms that allow seamless integration with soft tissue systems for such applications are urgently needed. Here, silver nanowires (Ag NWs)-based transparent microelectrodes and interconnects are designed to meet this demand. The Ag NWs percolating networks are patterned on flexible polymer substrates using an innovative photolithography-based solution-processing technique. The resulting nanowire networks exhibit a high average optical transparency of 76.1-90.0% over the visible spectrum, low normalized electrochemical impedance of 3.4-15 Ω cm^2^ at 1 kHz which is even better than those of opaque solid Ag films, superior sheet resistance of 11-25 Ω sq^−1^, excellent mechanical stability up to 10,000 bending cycles, good biocompatibility and chemical stability. Studies on Langendorff-perfused mouse and rat hearts demonstrate that the Ag NWs microelectrodes enable high-fidelity real-time monitoring of heart rhythm during co-localized optogenetic pacing and optical mapping with negligible light-induced electrical artifacts. This proof-of-concept work illustrates that the solution-processed, transparent, and flexible Ag NWs networks are a promising candidate for the next-generation of large-area multifunctional biointerfaces for interrogating complex biological systems in basic and translational research.

## 1. Introduction

Bioelectronic devices that provide multifunctional sensing and modulation of cell activity at the soft tissue interfaces represent powerful tools for many areas of biological research. Electrophysiology techniques have been the gold standard for investigating biological systems with high temporal resolution in basic research and clinical medicine for many decades.^[1,2]^ On the other hand, optophysiology techniques such as optogenetics and optical mapping can provide cell type-specific manipulation and mapping of cells or circuits with high spatial resolution, both *in vitro* and *in vivo* through the incorporation of light-sensitive proteins or utilizing intrinsic fluorescent reporters.^[3–5]^ Engineered platforms that allow multifunctional electrical and optical operations are of particular interest to combine the advantages of both techniques.

State-of-the-art electrophysiology tools rely on opaque metal microelectrodes to interface with bioelectric organ systems, such as the heart and brain.^[6–8]^ However, opaque microelectrodes are not ideal to bridge electrophysiology and optophysiology, because they not only produce significant light-induced electrical artifacts that scale with optical powers and distort recorded signals but also prevent light delivery to cells underneath the microelectrodes.^[9,10]^ Optically transparent microelectrodes based on indium tin oxide (ITO),^[11]^ graphene,^[12,13]^ carbon nanotubes (CNTs),^[14]^ metal nanomesh,^[15]^ and metal nanogrid^[16]^ have recently been reported to overcome those issues. Although flexible ITO/polyethylene terephthalate (PET) films are now commercially available for optoelectronic applications, their mechanical compliance and cyclic flexibility are inadequate for biological research due to the brittle nature of ITO. While graphene and CNTs present remarkable optical properties, their material quality is sensitive to fabrication methods.^[17]^ CNTs also suffer from controversial results and concerns associated with their cytotoxicity as biointerfaces.^[18–20]^ The sheet resistance (R_sh_) of graphene and CNTs is typically 1-2 orders of magnitude higher than that of ITO.^[12,14]^ As a result, graphene and CNTs microelectrodes require additional metal interconnects which complicates the device manufacturing process. To achieve a low electrochemical impedance, multiple graphene monolayers need to be stacked together while the gold nanomesh microelectrodes require extra conductive polymer coatings, which result in reduced optical transmittance.^[12,21]^ For gold nanogrid microelectrodes, electron-beam lithography is used for device fabrication, which imposes challenges for mass production.^[16]^

One-dimensional (1D) silver nanowires (Ag NWs) offer outstanding electrical conductivity, optical transparency, and mechanical flexibility. They have been widely used as flexible transparent electrodes in traditional electronic devices such as solar cells, displays, and light-emitting diodes (LEDs).^[22,23]^ The superior mechanical properties result from the intersliding behavior of the 1D Ag NWs. The open areas between the nanowires allow light to propagate with high optical transparency. Meanwhile, Ag NWs networks exhibit inherent high surface roughness because they are not strictly in the same film plane,^[22]^ which is a known factor to increase the effective interfacial contact area at the biotic/abiotic interface and boost the electrochemical performance.^[24,25]^ Although Ag is approved for use in many medical devices such as wound dressings^[26]^ and catheters^[27]^, current bio-related applications for Ag NWs have so far mainly focused on wearable sensors and do not fully exploit their unique advantages.^[28,29]^

This work reports the development and validation of transparent and flexible Ag NWs microelectrodes and interconnects for high-performance multifunctional electrophysiology and optophysiology. The Ag NWs networks are fabricated on polymer substrates using an innovative high-resolution (line width ~15 μm) photolithography-based solution-processing technique. The Ag NWs exhibit high average optical transparency of 76.1-90.0% over the visible spectrum, excellent R_sh_ of 11-25 Ω sq^−1^ that eliminates the need for additional metal interconnect layers, and low normalized electrochemical impedance of 3.4-15 Ω cm^2^ at 1 kHz that outperforms the opaque solid Ag films (10 Ω cm^2^). The high transmittance minimizes the light-induced electrical artifacts at intensity levels relevant to optophysiology studies. In addition, the Ag NWs networks exhibit robust adhesion to the substrates and excellent mechanical flexibility with stable R_sh_ after 10,000 repetitive bending against a small radius of 5 mm, achieving a conformal contact with the soft and curvilinear tissue. *In vivo* histology studies reveal that the Ag NWs networks are biocompatible. *Ex vivo* electrophysiological recordings of electrogram (EG) signals from Langendorff-perfused mouse and rat hearts during cardiac dysfunction, optogenetic pacing, and optical mapping demonstrate that the Ag NWs microelectrodes and interconnects can be an enabling platform for studying the dynamics and operating principles of complex biological systems.

## 2. Results and Discussion

### 2.1. Design and Fabrication of Ag NWs Microelectrodes and Interconnects

Figure 1a demonstrates the schematic illustration of the simplified fabrication process. The process begins with laminating a thin PET film on the handling glass. Next, a 0.5 μm transparent SU-8 adhesive layer is coated on the PET film, soft baked and exposed by ultraviolet (UV) illumination. Ag NWs solutions are then uniformly spin coated on SU-8. After that, the Ag NWs/SU-8 mixtures are hard baked to allow crosslinking of UV exposed SU-8. During this step, Ag NWs are partially embedded into the SU-8 layer to improve adhesion with the PET substrate. Afterwards, photolithographic patterning and wet etching define the Ag NWs microelectrodes and interconnects. A 7 μm SU-8 layer encapsulates the device before it is released from the handling glass. This fabrication strategy contrasts sharply to conventional metal,[15,30] CNTs,[14,31] and graphene microelectrodes,[32,33] which require costly high vacuum deposition tools and/or multistep growth and transfer processes. The fabrication details are described in the Experimental Section.

**Figure 1.**
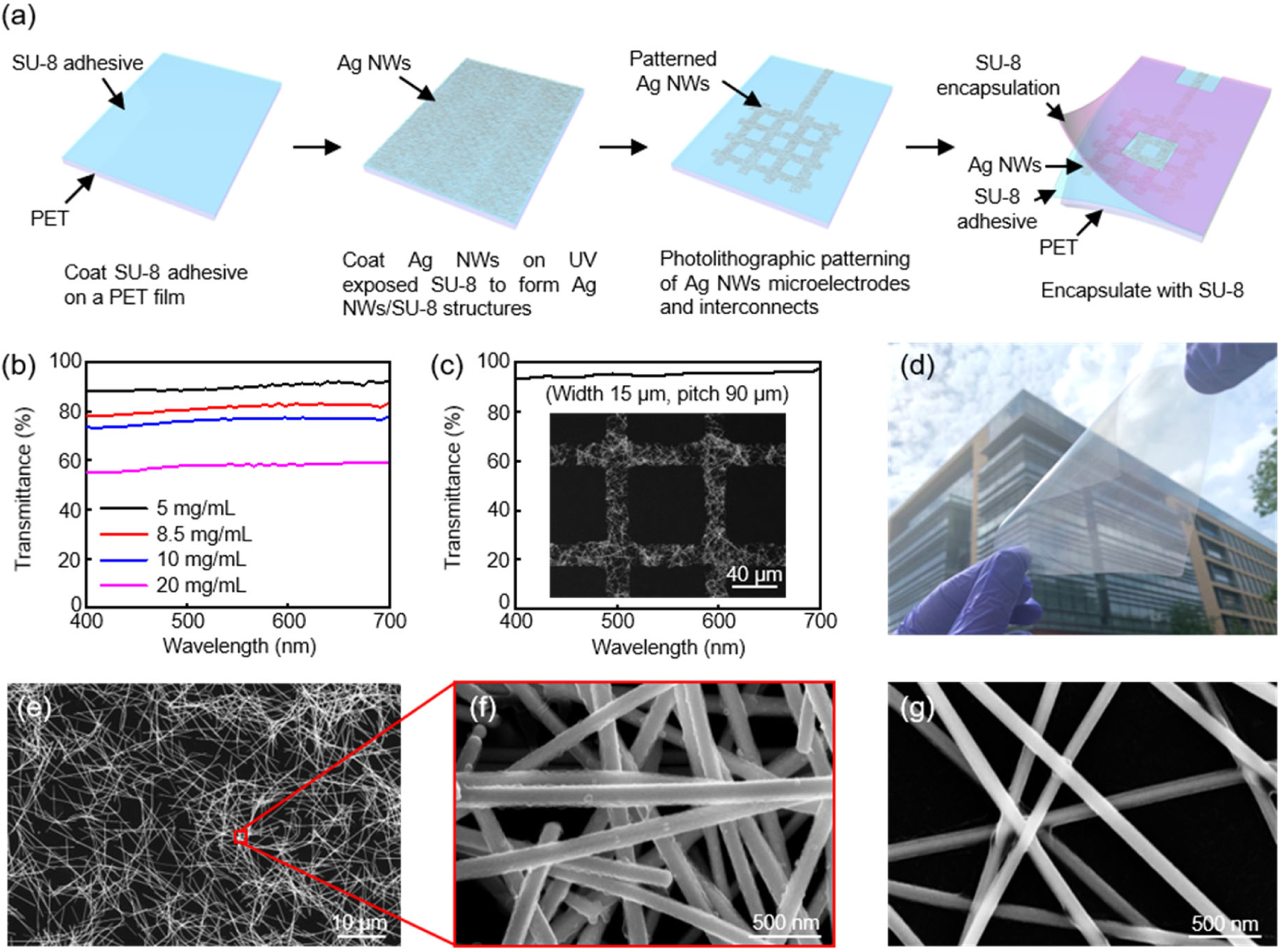
a) Schematic illustration of the fabrication steps for the flexible and transparent Ag NWs microelectrodes and interconnects. b) Transmission spectra of Ag NWs networks at various concentrations. c) Transmission spectrum of a patterned Ag NWs grid. (Inset) SEM image of the grid patterns. d) Optical image of a 10 × 10 cm^2^ Ag NWs/SU-8/PET film. SEM images of Ag NWs at e-f) 8.5 mg/mL, and g) 5 mg/mL, respectively.

### 2.2. Optical, Electrical, and Electrochemical Properties of Ag NWs Microelectrodes and Interconnects

The NWs network density is a crucial parameter that affects the physical properties of the resulting microelectrodes and interconnects. Decreasing the network density will increase the open areas between the NWs to achieve high optical transparency. Meanwhile, this will reduce the conductive electrical pathways and effective interfacial areas within the same geometrical surface areas, resulting in decreased electrical and electrochemical performance. In this work, the Ag NWs network density is controlled by changing the concentrations of Ag NWs solutions used in spin coating. The average transmittance of Ag NWs networks between 400 and 700 nm increases from 57.7% to 76.1%, 81.3%, and finally 90.0% with the Ag NWs concentrations decreasing from 20 to 10, 8.5, and 5.0 mg/mL, respectively (Figure 1b). Importantly, the photolithographic fabrication process allows preparation of the conductive Ag NWs networks with a resolution down to 15 μm, which is one of the highest resolutions among the reported patterning technologies for conductive Ag NWs structures.^[34–37]^ Figure 1c shows a patterned Ag NWs grid structure (a width of 15 μm and a pitch of 90 μm) with a superior average transmittance of 95.2%. In addition, the fabrication strategy enables the scaling up of the Ag NWs structures for large-area biointerfacing. Figure 1d displays a 10 × 10 cm^2^ Ag NWs/SU-8/PET film with high transparency and uniform dispersion of Ag NWs networks. The Ag NWs have a diameter of ~120 nm and a length between 10 and 20 μm, as observed in the scanning electron microscope (SEM) image in Figure 1e. As expected, decreasing the Ag NWs concentrations leads to a less dense network (Figure 1e,f).

Figure 2a depicts the R_sh_ versus transmittance of the Ag NWs/SU-8/PET films. The Ag NWs interconnects exhibit R_sh_ values in the range of 4.1-25 Ω sq^−1^ with optical transmittance ranging between 57.7% and 90.0%, respectively. Further increasing the transmittance to 93.0% rapidly increases R_sh_ to 80 Ω sq^−1^ due to the dramatically reduced conductive electrical pathways. The R_sh_ is comparable to rigid ITO^[38]^ and outperforms carbon-based flexible transparent electrodes such as CNTs^[14]^ and graphene^[12]^ at similar optical transparency levels. The superior electrical performance allows the Ag NWs to serve as the conductive interconnects for the microelectrodes.

**Figure 2.**
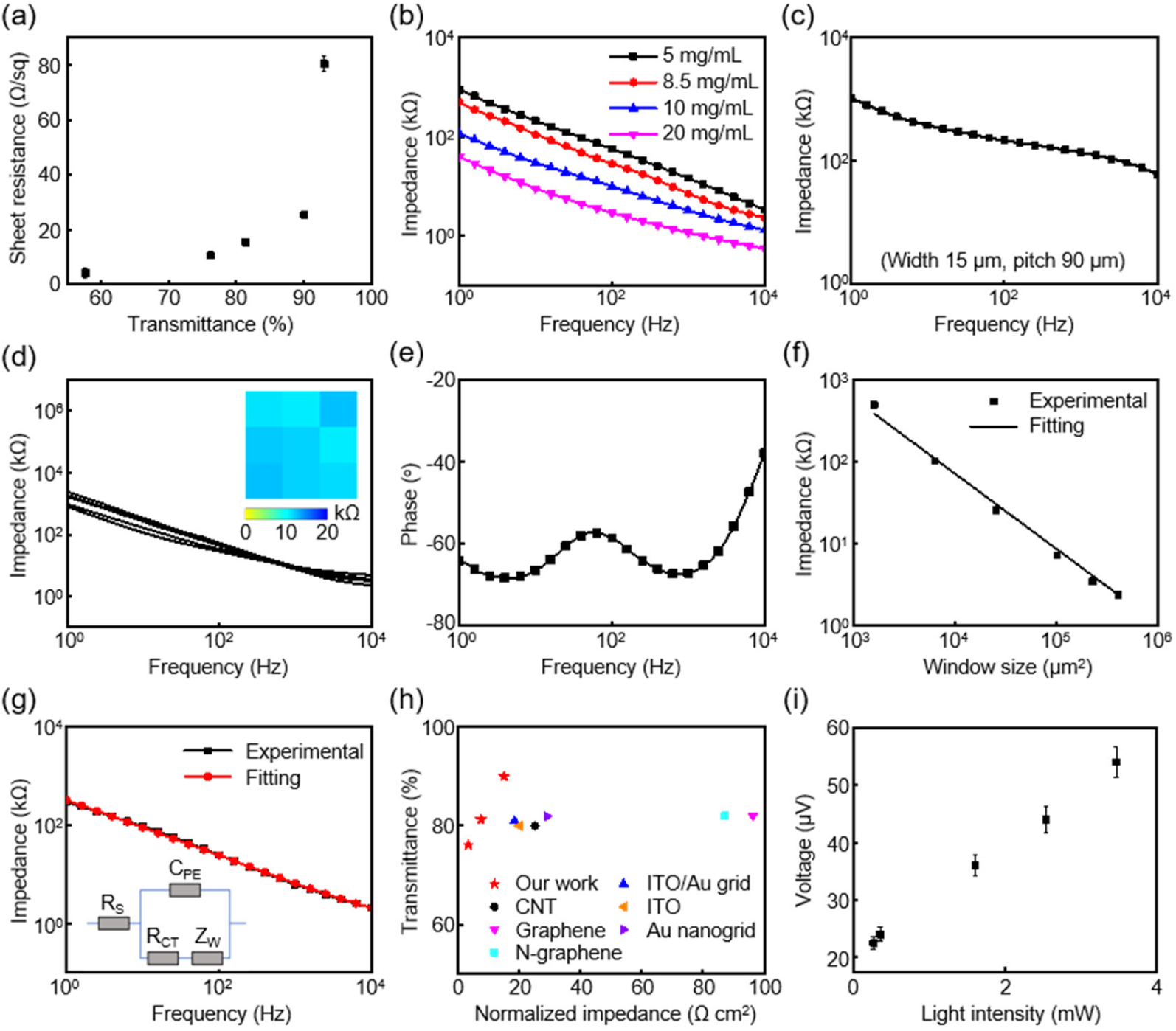
a) Sheet resistance versus transmittance of the Ag NWs interconnects. b) Impedance plots of Ag NWs microelectrodes at various concentrations. The microelectrode size is 320 × 320 μm^2^. c) Impedance plot of the grid patterned Ag NWs microelectrodes. d) Impedance spectra of 9 microelectrodes in a 9-channel Ag NWs microelectrode array. e) Phase plot of the Ag NWs microelectrodes. f) Impedance of the Ag NWs microelectrodes as a function of microelectrode size. g) EIS results and equivalent circuit model fitting. h) Normalized impedance of Ag NWs microelectrodes versus transmittance. Results are compared to major reported microelectrodes with transmittance over 80%. i) Light-induced electrical artifacts of the Ag NWs microelectrodes.

Electrochemical impedance is one of the most important characteristics of a microelectrode. A low impedance improves the recording quality with a high signal-to-noise ratio (SNR). Figure 2b presents the impedance of Ag NWs microelectrodes with frequencies ranging from 1 Hz to 10 kHz in a phosphate-buffered saline (PBS) solution measured by electrochemical impedance spectroscopy (EIS). The microelectrode dimension is 320 × 320 μm2, if not specifically mentioned. Impedance at 1 kHz is widely used for comparison among different microelectrodes. The impedance values at 1 kHz for the Ag NWs microelectrodes decrease from 14.7, to 7.2, 3.3, and 1.2 kΩ with Ag NWs concentrations increasing from 5.0, to 8.5, 10, and 20 mg/mL, respectively. This is because the effective interfacial areas increase with higher Ag NWs concentrations. The patterned Ag NWs grid microelectrodes (transmittance of 95.2%) in Figure 1c possess a moderate impedance of 137.2 kΩ at 1 kHz (Figure 2c), providing a good candidate for electrophysiological applications requiring extremely high optical transparency. Figure 2d shows the impedance response from 9 microelectrodes in a 9-channel array (3 × 3, pitch of 2.5 mm) with the Ag NWs concentration of 8.5 mg/mL. The microelectrode array (MEA) exhibits uniform performance with an average impedance of 8.3 ± 0.6 kΩ. The low impedance and excellent uniformity are crucial for large-scale and high-fidelity electrical recording. The phase responses in Figure 2e indicate the Ag NWs are highly capacitive (phase angle between −70o and −60o) at physiologically relevant low frequencies (1 Hz to 1 kHz) and become more resistive at higher frequencies. Figure 2f illustrates the impedance of the Ag NWs microelectrodes at 1 kHz as a function of microelectrode size. Clearly, the impedance demonstrates a near-precise dependance on the microelectrode area, suggesting a capacitive interface.^[21]^

To better understand the electrochemical properties of the Ag NWs microelectrodes, the EIS measurement results are fit to an equivalent circuit model in Figure 2g.^[15,32]^ The circuit model consists of solution resistance (R_S_), constant phase element (C_PE_), charge transfer resistance (R_CT_) and Warburg element for diffusion (Z_W_). C_PE_ is defined by 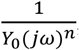, where *Y*_*0*_ is the magnitude of C_PE_, *j* is the unit imaginary number, *ω* is the angular frequency, and *n* is a constant accounting for the character of capacitance. The n values range from 0 to 1, where n = 0 represents a pure resistor and n = 1 represents an ideal capacitor. The model provides a good fit for the Ag NWs microelectrodes, as plotted in Figure 2g. Table 1 summarizes the fitting parameters. The n value (0.7) represents a more capacitive interface.

**Table 1.** Equivalent circuit fitting results of Ag NWs microelectrodes.

| $R_s (\Omega)$ | $C_{PE} (S \times s^n)$ | | $R_{CT} (\Omega)$ | $Z_W (S \times s^{1/2})$ |
| --- | --- | --- | --- | --- |
|  | Q | n |  |  |
| 830.0 | $2.1 \times 10^{-6}$ | 0.7 | $1.3 \times 10^3$ | $9.5 \times 10^{-6}$ |

The electrochemical and optical performance of Ag NWs microelectrodes are compared to previously reported state-of-the-art transparent microelectrodes for electrophysiological studies with a transmittance >80.0%, including CNTs,^[14]^ graphene,^[13]^ nitrogen-doped graphene (N-graphene),^[39]^ ITO,^[40]^ Au nanogrid,^[16]^ and ITO/Au grid^[41]^ (Figure 2h). All impedance values at 1 kHz are normalized to the areas of the microelectrodes for fair comparison. The Ag NWs microelectrodes show both low normalized impedance (7.4 to 15 Ω cm^2^) and high optical transmittance (81.3% to 90.0%), which are among the most competitive performance among transparent microelectrodes for biointerfacing. The normalized impedance of Ag NWs microelectrodes (7.4 Ω cm^2^) at 81.3% transmittance is even better than those from opaque solid Ag film microelectrodes (10 Ω cm^2^) with a thickness (120 nm) similar to the diameter of the Ag NWs (Figure S1, Supporting Information). The root-mean-square (RMS) roughness of the Ag NWs and Ag films determined from atomic force microscopy (AFM) measurements is 130 and 6.7 nm, respectively (Figure S2, Supporting Information). Compared with planar microelectrodes, the Ag NWs microelectrodes here possess significantly larger effective electrode/electrolyte interfacial areas due to higher surface roughness, leading to better electrochemical performance. In view of the balanced electrical, electrochemical, and optical properties, the Ag NWs networks with 81.3% transmittance are used in subsequent experiments.

Light-induced electrical artifacts of the Ag NWs microelectrodes are measured in a modified Tyrode’s solution with a blue microscale LED placed next to the microelectrodes as the light source. The artifacts increase linearly with light intensity (Figure 2i). The amplitude is below 60 μV under a 10 ms pulsed blue light (462 nm) illumination with an intensity of 3.5 mW, relevant to optophysiology studies.[42] The small artifacts are due to the high optical transparency of the microelectrodes, limiting the amount of light absorbed. Those values are much smaller than typical local field potentials (0.5 to 5 mV).^[43]^

Figure 3a shows the benchtop recording results of a 10 Hz sine wave signal with 20 mV peak-to-peak amplitude in a PBS solution using the Ag NWs microelectrodes. No decrease in signal amplitude is observed, suggesting a high-fidelity recording performance. Figure 3b presents the power spectrum density (PSD) of the recorded signals to provide more details on noise and signal in the frequency domain. The large peak at 10 Hz is from the input signal. The Ag NWs microelectrodes show an RMS noise and a SNR of 48.3 μV and 42.7 dB, respectively. To evaluate the chronic chemical stability of the Ag NWs microelectrodes, the impedance is continuously measured over 1 month with the microelectrodes immersed in a PBS solution at 37 °C. Oxidation and corrosion of Ag will decrease the effective interfacial area and increase the impedance. As shown in Figure 3c, the microelectrodes exhibit stable electrochemical performance, showing no corrosion of Ag in the PBS solution for 1 month and a good adhesion among Ag NWs, SU-8, and PET without delamination. Figure 3d shows cyclic voltammetry (CV) results of the Ag NWs microelectrodes at a scan rate from 50 to 200 mV s^−1^ in a PBS solution. The voltage ranges from −0.8 to 0.8 V. The Ag NWs microelectrodes exhibit a CV morphology in agreement with previously reported electrochemical properties of Ag electrodes.^[44,45]^ The peak current shows a linear dependence on the square root of scan rate (Figure S3, Supporting Information), indicating that charge transfer at the Ag NWs/electrolyte interface is diffusion controlled.^[46]^

**Figure 3.**
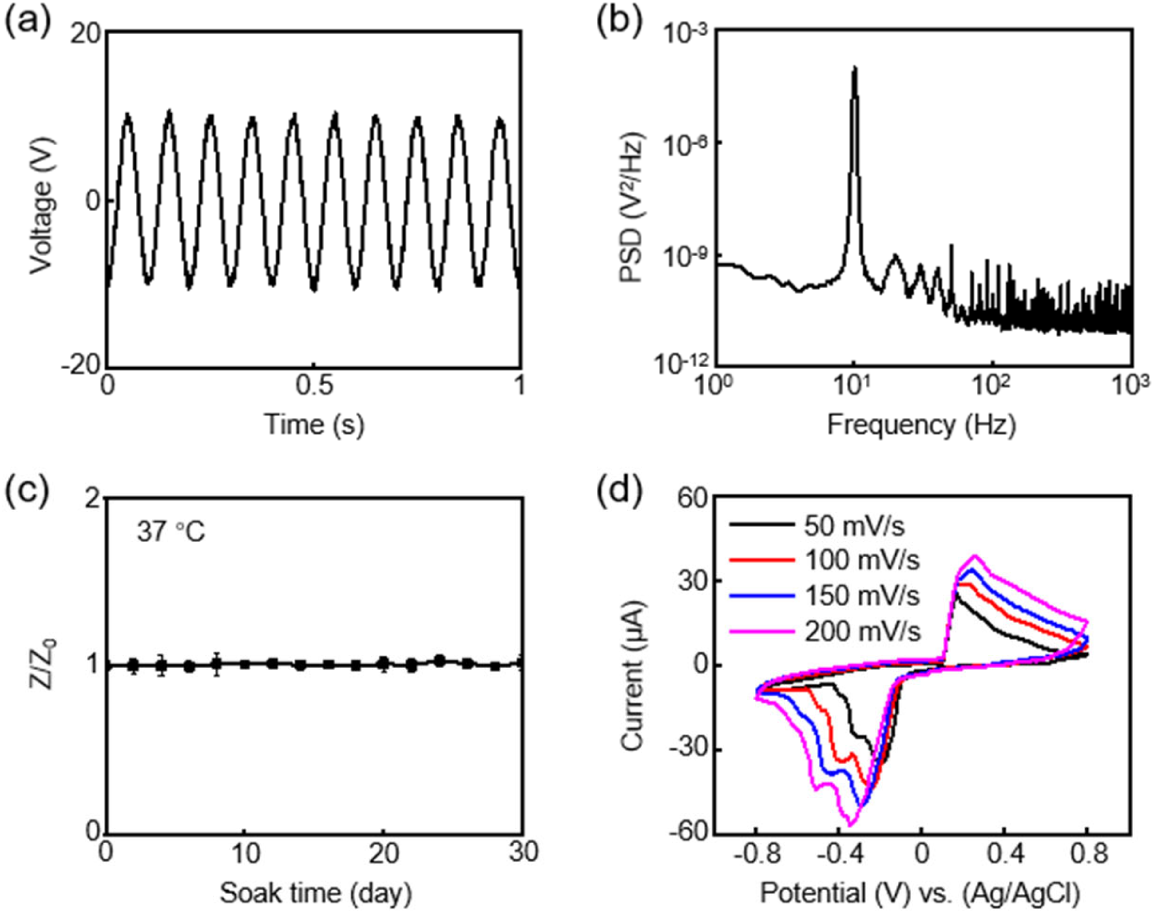
a) Electrical benchtop recording output from the Ag NWs microelectrodes, with 10 Hz, 20 mV peak-to-peak sine wave input. b) PSD of the signals recorded by the Ag NWs microelectrodes in (a). c) Soak test of the Ag NWs microelectrodes in a PBS solution at 37 °C. Z0 is the impedance at day 0, whereas Z represents the impedance at a specific day. d) CV curves of the Ag NWs microelectrodes at various scan rates.

### 2.3. Mechanical Properties of Ag NWs Microelectrodes and Interconnects

Mechanical flexibility of a bioelectronic device is important to achieve conformal contact at the biotic/abiotic interface and sufficient mechanical compliance to various external deformations. Figure 4a presents the mechanical bending stability of the Ag NWs/SU-8/PET film against a 0.5 cm bending radius. This radius is similar to the anatomy of interest in small animal models.^[13,47]^ The result shows negligible change in R_sh_ even after 10,000 bending cycles. The mechanical stability of the Ag NWs networks is improved compared to the Ag film (thickness 120 nm) which shows a 117 times increase in R_sh_ after 5,000 cycles. Thus, the Ag NWs structures are suitable for reliable biointerfacing and repetitive use. Figure 4b,c demonstrate optical images of a blue LED connected to a voltage supply using the Ag NWs interconnects. The resistance of the system stays unchanged under 10,000 bends of the interconnects, which is known from the current and voltage readings shown on the voltage supply.

**Figure 4.**
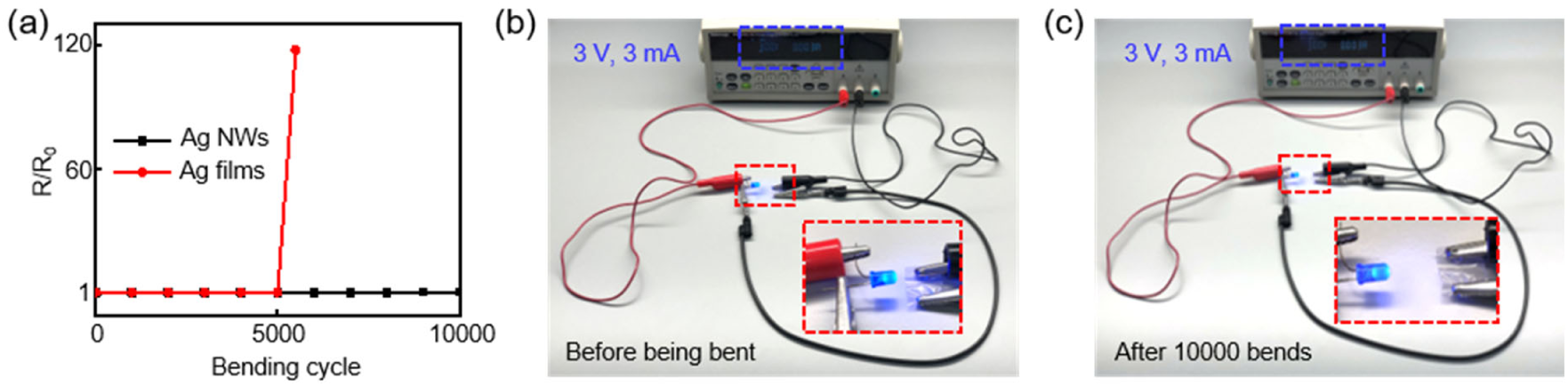
a) Variation of sheet resistance versus bending cycle for Ag NWs interconnects and solid Ag films at a bending radius of 0.5 cm. R0 is the sheet resistance before being bent, whereas R represents the sheet resistance at a specific bending cycle. Optical images of an operating blue LED connected by Ag NWs/SU-8/PET films b) before being bent, and c) after 10,000 cycles at a bending radius of 0.5 cm.

### 2.4. *Ex Vivo* Cardiac Electrical Recording with Optogenetic Pacing

Recording and modulating EG signals of transgenic mouse hearts expressing one of the most widely used blue light excitable opsins, channelrhodopsin-2 (ChR2)^[48,49]^, provides proof-of-concept demonstrations to evaluate the general utility of the Ag NWs microelectrodes and interconnects for electrophysiology and optogenetics studies. Figure 5a shows the experimental setup, where the highly flexible microelectrodes conformally contact the right ventricle of the Langendorff-perfused mouse heart. Photons from an external blue microscale LED (462 nm and 1.5 mW) pass through the transparent Ag NWs microelectrodes to pace the underneath cardiomyocytes. Standard commercial far-field electrocardiogram (ECG) electrodes serve as the reference. Atrioventricular (AV) block, a common heart rhythm disorder, is chosen as the disease model in this study. AV block is introduced by ischemic low perfusion flow.^[16]^ Figure 5b shows the baseline electrical recording results using the commercial reference and Ag NWs microelectrodes without optogenetic pacing. AV block is confirmed by the skipped beat (i.e., dissociation between P wave and QRS complex) in the baseline signals. Importantly, the morphology of the EG signals recorded from Ag NWs microelectrodes is almost identical to the far-field ECG recorded from the commercial reference. The average durations of five QRS complexes from the Ag NWs microelectrodes and the reference electrode are 9.7 ± 0.5 ms and 9.8 ± 0.4 ms, respectively. Those results indicate the high-fidelity recording capabilities of the Ag NWs microelectrodes.

**Figure 5.**
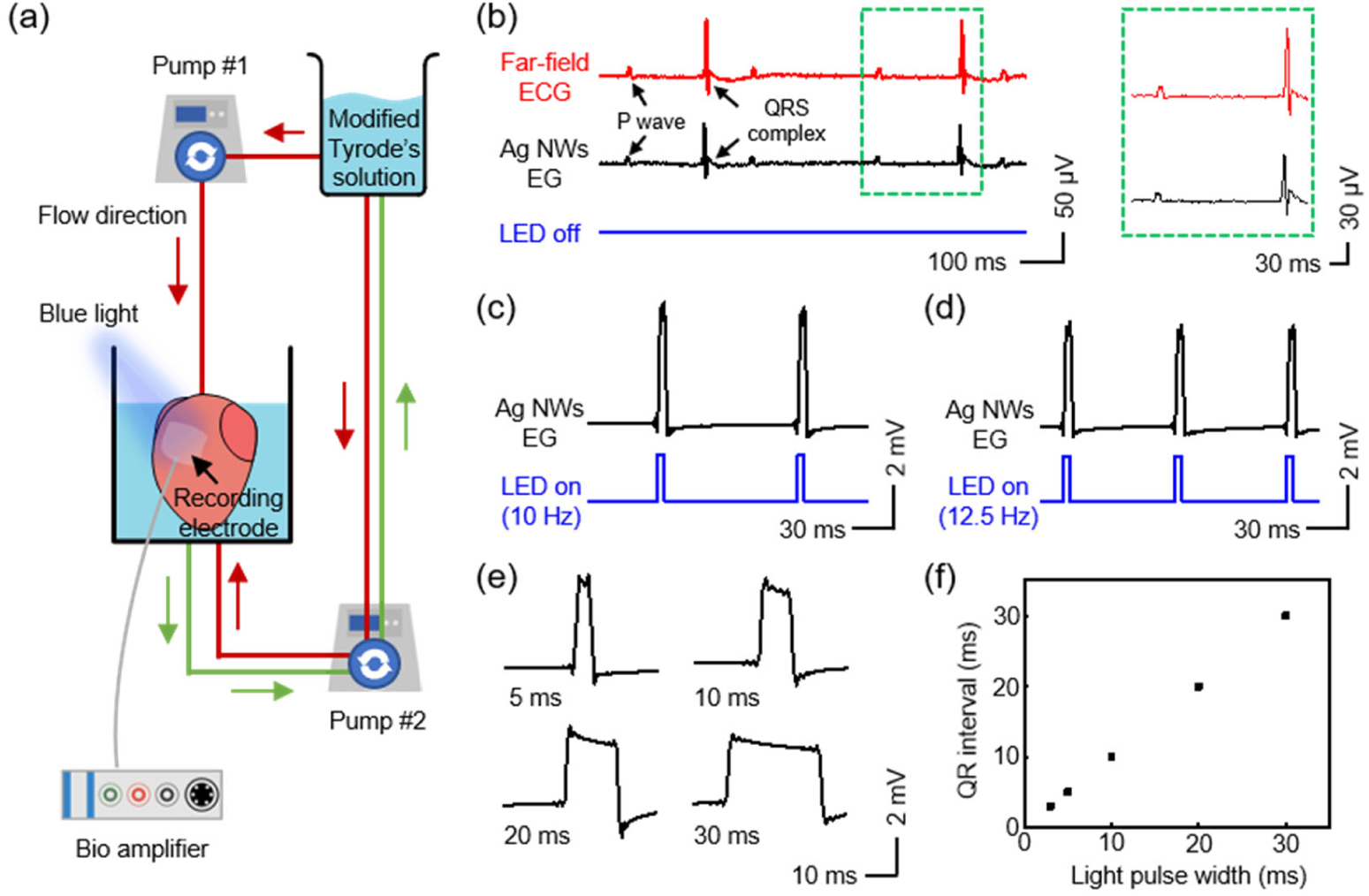
a) Schematic of the Langendorff perfusion experimental setup for electrical recording and optogenetic pacing of a ChR2-expressing mouse heart. b) (Left) Electrical recordings of AV block from the reference electrode and Ag NWs microelectrodes. (Right) Detailed representative P wave and QRS complex recorded by the reference electrode and Ag NWs microelectrodes. EG recording from the Ag NWs microelectrodes during co-localized optogenetic pacing at c) 10 Hz and d) 12.5 Hz. e) Representative QRS complexes recorded by the Ag NWs microelectrodes at different optogenetic pacing pulse widths. f) QR interval recorded by the Ag NWs microelectrodes versus light pulse width for optogenetic pacing.

Optogenetic pacing of the cardiomyocytes expressing ChR2 underneath the Ag NWs microelectrodes at a frequency higher than the intrinsic heart rhythm is then performed to treat AV block and restore the normal sinus rhythm of the heart. The intrinsic heart rhythm of the mouse heart with 2^nd^ degree AV block in Figure 5b is ~240 beats per minute (4 Hz). Figure 5c,d present the recorded EG signals from the Ag NWs microelectrodes during 5 ms optogenetic pacing at 10 and 12.5 Hz with duty cycles at 5% and 6.25% (blue curves), respectively. Clearly, the Ag NWs microelectrodes allow efficient light delivery to activate the cardiomyocytes under the microelectrodes and capture the resulting heart rhythm. The amplitude of the EG (5 mV) from the Ag NWs microelectrodes is comparable to that of the far-field ECG signals from the reference electrode that is not directly exposed to blue light illumination (Figure S4, Supporting Information). In addition, the signal amplitude is over two orders of magnitude higher than the light-induced electrical artifacts in Figure 2i. Together, those results suggest the Ag NWs microelectrodes enable light-induced electrical artifacts-free recordings. Figure 5e shows the recorded QRS complexes by the Ag NWs microelectrodes with 10 Hz optogenetic pacing at various pulse width (5.0, 10, 20, 30 ms). The duration of the QRS complexes (7.2, 13, 23, 33 ms) is accurately controlled by the pulse width. In addition, the QR intervals directly corresponds to the applied pulse durations (Figure 5f).

### 2.5. *Ex Vivo* Cardiac Electrical Recording with Optical Mapping

Voltage mapping on Langendorff-perfused rat hearts under electrical pacing by a complementary metal-oxide-semiconductor (CMOS) imaging system is used to validate the feasibility of optical mapping through the transparent Ag NWs microelectrodes. An adult rat heart is stained with the voltage-sensitive fluorescent dye di-4-ANEPPS. A flexible 3 × 3 Ag NWs MEA (pitch 2.5 mm) forms a conformal and intimate contact to the anterior to lateral side of the left ventricle. The fluorescent dye is excited at 530 nm with a maximum fluorescence emission at 705 nm, filtered with a 600 nm long-pass filter. The Ag NWs MEA exhibits high transmittance values of 81.4% and 82.4% at the two wavelengths to allow photons to pass for efficient co-localized optical mapping, respectively. This is confirmed by the representative time aligned EG signals and optical fluorescence measured from the same location of the left ventricular surface under external electrical pacing in Figure 6a. The electrical pulses are delivered by a platinum pacing electrode with 2 ms pulse duration and twice the amplitude of the pacing threshold. The representative optical and electrical activation maps at a 5 Hz electrical pacing frequency show strong correlations (Figure 6b,c), where the optical signals represent average results over ~0.5 to 1 mm of depth below the epicardial surface of the heart tissue,^[50]^ and the Ag NWs MEA records EG signals directly from the tissue surface. The above results suggest that the Ag NWs MEA enables accurate co-localized optical and electrical mapping of heart rhythms.

**Figure 6.**
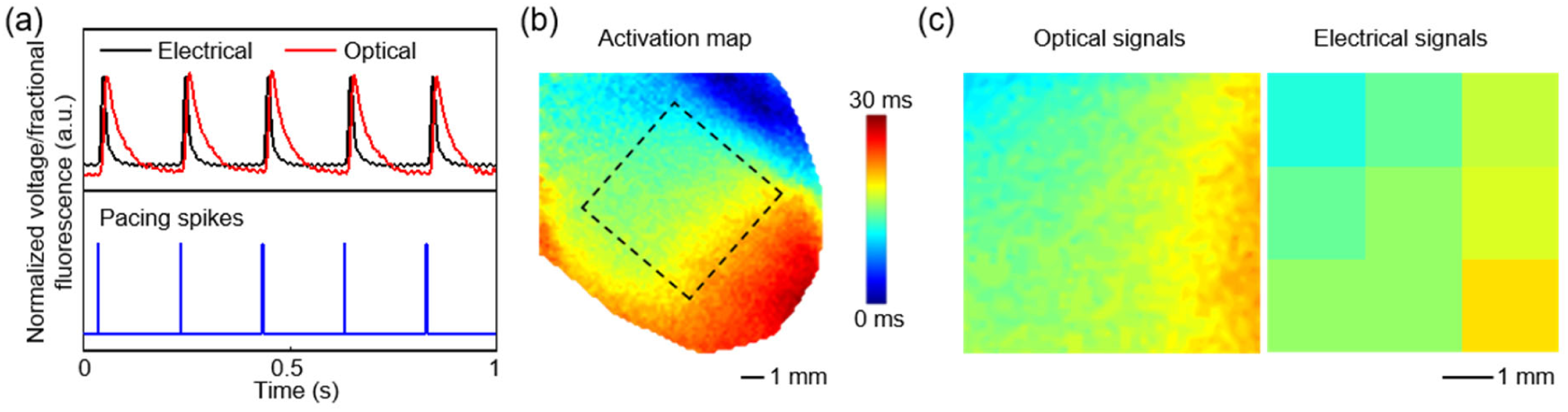
a) Representative optical and electrical signals recorded at the same sites on a Langendorff-perfused rat heart under electrical pacing at 5 Hz (a.u., arbitrary units). b) Optical activation map from the left ventricle of the rat heart. The black dashed square shows the position of the Ag NWs MEA. c) Activation maps obtained from the optical and electrical data for the region under the Ag NWs MEA.

### 2.6. Histological Evaluation of Biocompatibility

To evaluate the *in vivo* biocompatibility of the devices, the Ag NWs microelectrodes are implanted onto the anterior epicardial surface of the ventricles in rats. Four weeks after implantation, the ventricular tissues are collected, fixed, and stained with Masson’s trichrome where blue represents fibrotic tissue, pink represents myocytes, and white represents interstitial space, respectively (Figure 7a,b). Compared with the healthy myocardium, there is an increase in fibrosis at the site where the device is attached to the heart. This response is expected due to inflammation caused by a foreign body response. Importantly, there is no significant difference in the percent volume of myocytes, fibrosis, and interstitial space in the transmural cross section of the heart between control animals and those implanted with devices (p<0.05, Figure 7c). Overall, the Ag NWs microelectrodes are reasonably biocompatible with the heart and not causing major complications. In the future studies we will employ anti-inflammatory strategies to reduce the inflammation and fibrosis.

**Figure 7.**
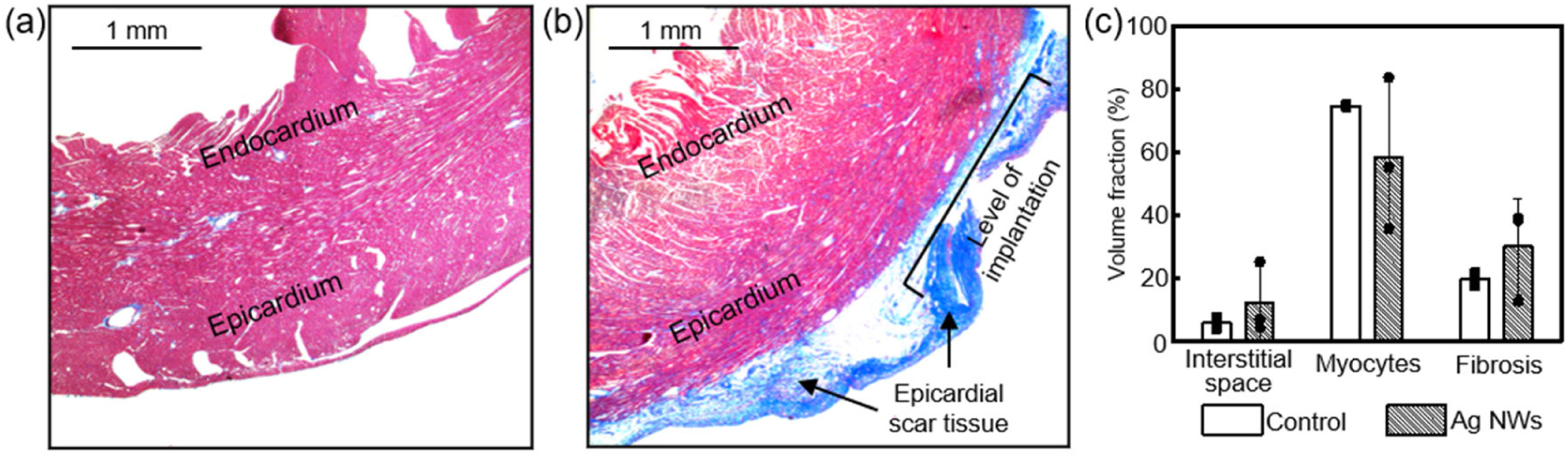
*In vivo* evaluations of biocompatibility of the devices. Representative images of Masson’s trichrome-stained cross sections of the ventricular tissue near the site of device attachment of a) a rat with no device implanted and b) a rat with device implanted. With this Masson’s trichrome stain, pink represents cardiomyocytes, blue represents fibrotic tissue, and white represents interstitial space. c) Quantification of the cardiomyocytes, fibrotic tissue, and interstitial space in the cross sectional epicardium of the heart (n = 3 independent animals per condition). No statistically significant differences were observed in the volume fraction composition of the myocardium between the control animals and those implanted with devices (p<0.05).

## 3. Conclusion

In this work, Ag NWs-based flexible and transparent microelectrodes and interconnects are successfully demonstrated. The Ag NWs interfaces are developed by a simple high-resolution photolithography-based solution-processing technique that is compatible with large-area fabrication. The resultant Ag NWs networks exhibit superior optical, electrical, electrochemical, and mechanical performance, excellent biocompatibility and chemical stability for biointerfacing. Benchtop characterizations reveal that the integration schemes can achieve MEAs with high uniformity. Proof-of-concept demonstrations show that the Ag NWs microelectrodes allow high-fidelity co-localized monitoring of heart rhythms during optogenetic pacing and optical mapping with negligible light-induced electrical artifacts. This study expands the biomedical applications of metal NWs and shows that Ag NWs microelectrodes and interconnects are promising candidates for the next generation of multifunctional biointerfaces combining electrophysiology with optophysiology to measure and modulate bioelectric organ systems.

## 4. Experimental Section

### Microelectrodes and Interconnects Fabrication

Firstly, a PET film (25 μm thick, CS Hyde Company) was laminated on a glass slide using polydimethylsiloxane as the adhesive. A 0.5 μm thick SU-8 photoepoxy (SU-8 2000.5, Microchem) was then spin coated on the PET film, followed by soft baking at 65 °C and 95 °C for 2 minutes each. Ag NWs/isopropyl alcohol solutions (ACS Material) at different concentrations were spin coated on the SU-8 layer after UV exposure. The mixtures were then hard baked at 95 °C and 110 °C for 3 and 15 minutes, respectively. To define the Ag NWs patterns, a photoresist (AZ P4620, Integrated Micro Materials) was used as a hard mask, followed by wet etching in silver etchants. Finally, the device was encapsulated by a 7 μm thick SU-8 photoepoxy (SU-8 2007, Microchem) via photolithography.

### Optical Measurement

A spectrophotometer (V-770 UV-vis/NIR, Jasco Inc.) was used to measure transmission spectra of the Ag NWs films. A spectrometer (AvaSpec-ULS2048L, Avantes) measured the emission profiles of the LEDs.

### SEM Characterization

The morphology of the Ag NWs networks was studied by SEM (PIONEER EBL, Raith Inc.) at an accelerating voltage of 10.0 kV.

### Electrical Measurement

A four-point probe (SRM-232, Guardian Manufacturing Inc.) was used to measure the R_sh_ of Ag NWs networks.

### Electrochemical Measurement

A Gamry potentiostat (Reference 600+, Gamry Instruments Inc.) was used to conduct electrochemical measurements, including EIS and CV. EIS tests were performed at a frequency ranging from 1 Hz to 10 kHz. CV tests were measured within a potential window from −0.8 to 0.8 V. The electrochemical measurements used a three-electrode configuration in a 1 × PBS solution (Sigma-Aldrich), where the Ag NWs microelectrodes, a platinum electrode, and an Ag/AgCl electrode served as the working, counter, and reference electrodes, respectively.

### AFM Measurement

The roughness of Ag NWs and solid Ag films was measured with a MultiMode 8-HR AFM (Bruker Inc.) using a ScanAsyst - Air probe.

### Electrical Benchtop Test

A PowerLab data acquisition system (PowerLab 16/35, ADInstruments Inc.) was used to deliver a 10 Hz sine wave signal with 20 mV peak-to-peak amplitude through a platinum electrode into a 1 × PBS solution (Sigma-Aldrich). The microelectrodes and another platinum electrode were connected to the system at different channels for comparison. MATLAB was used to process the PSD signals.

### Mouse Model

For optogenetics, mice that express ChR2 in cardiomyocytes alone were created by cross-breeding one parent (Stock No. 011038, Jackson Labs)^[51]^ with another parent (Stock No. 024109, Jackson Labs)^[52]^.

### Animal Experiments

All animal procedures were performed according to protocols approved by the Institutional Animal Care and Use Committee of The George Washington University and in conformance Guide for the Care and Use of Laboratory Animals published by the National Institutes of Health.

For *ex vivo* optogenetics and electrical recording demonstrations, adult mice expressing ChR2 in the cardiomyocytes were used. Mice were anesthetized by inhaling a mixture of 5% isoflurane vapors with 2 mL/minute oxygen flow in an induction chamber for 3-5 minutes. Cervical dislocation was immediately performed after toe pinch verification of loss of pain sensation. The hearts were quickly excised after thoracotomy, cannulated via the aorta, and placed into a Langendorff-perfused constant flow system with a modified Tyrode’s solution (in mм: NaCl 140, KCl 4.7, MgCl2 1.05, CaCl2 1.3, HEPES 10, Glucose 11.1, BDM 15, pH 7.4 at 37 °C) bubbled with 100% O2. Hydrostatic pressure was maintained at 70 ± 5 mmHg. Electrical recordings were conducted in a three-electrode configuration with Ag NWs microelectrodes or far-field needle electrodes (MLA1203, ADInstruments Inc.) as the working electrode. A blue LED was placed directly on top of the Ag NWs microelectrodes which were in contact with the posterior side of the hearts. Cardiomyocytes were activated by photons passing through the transparent microelectrodes. Optical stimulation was set at 10 Hz (pulse width of 3, 5, 10, 20, 30 ms) and 12.5 Hz (pulse width of 5 ms) where the rate of stimulation is faster than the intrinsic heart rate of the heart. Pacing and capture was verified in both reference EG and Ag NWs microelectrode recordings. EG signals were acquired at 1 kHz sampling frequency (0.5 Hz to 2 kHz bandpass filtered) and analyzed with the LabChart software (ADInstruments Inc.). AV block was induced by ischemia-reperfusion of the hearts. The total length of the experiment was within 3 hours.

For *ex vivo* synchronized optical and electrical mapping, an adult Sprague-Dawley female rat was deeply anesthetized with 5% isoflurane vapors at 2 mL/minute flow of O2 and humanely sacrificed by exvision of the heart and exsanguination. The excised heart was perfused in the constant pressure (70-80 mmHg) Langendorff-mode with the modified Tyrode’s solution bubbled with 100% O2. The heart was allowed to stabilize for about 10 minutes. The Ag NWs 9-channel MEA was placed on the anterior to lateral side of the left ventricle. A bipolar platinum pacing electrode was placed at the base of the left ventricle. The voltage-sensitive dye di-4-ANEPPS (Biotium, Catalog number #61010) was loaded into the heart via the coronary circulation. First, 20 μL of di-4-ANEPPS stock solution (1.25 mg/mL in dimethyl sulfoxide) was reconstituted in 1 mL of the modified Tyrode’s solution. Then 0.1 mL of the dye was slowly injected through the aortic cannula over 5 minutes. Any excess dye was washed out after dye loading. A LEX3-G Green LED System (peak wavelength at 530 nm, SciMedia, USA) with an excitation-filter (520 ± 17 nm) was used for excitation of di-4-ANEPPS. Fluorescence signals were filtered using a long-pass filter (600 nm) and collected by the Ultima-L CMOS camera system (SciMedia, USA) at a sampling frequency of 2000 Hz. A 15 × 15 mm2 field of view was projected to the 100 × 100 pixel CMOS sensor. The synchronized recording was conducted at a pacing cycle length of 200 ms, with the pacing stimulus set at twice the amplitude of the pacing threshold and 2ms pulse duration.

For *in vivo* histology experiments, male and female adult Sprague-Dawley and Long Evans rats were used for implantation (n = 3). All procedures were performed under general anesthesia administered by inhaled isoflurane vapors. Ventilation was provided by a VentElite small animal ventilator (Harvard Apparatus). Following exposure of the heart via left thoracotomy, the Ag NWs microelectrodes were sutured onto the anterior epicardial surface of the ventricles with 6-0 non-resorbable monofilament sutures. The thoracic cavity, muscle, and skin were closed in subsequent order. Appropriate post-operative care was provided, including analgesic doses of 0.05-0.1 mg/kg buprenorphine delivered via subcutaneous injection every 12 hours for up to 48 hours after surgery. 4 weeks after Ag NWs implantation, hearts were collected for histological analysis. Rats were sacrificed by inhalation of 5% isoflurane vapors at 2 mL/minute oxygen flow with an EZ anesthesia machine (EZ Systems Inc.). Following confirmation of cessation of pain via toe pinch, the hearts were excised. The aorta was cannulated and retrograde-perfused with 10% neutral-buffered formalin. Cross sections of the hearts were paraffin embedded, sectioned, and stained with Masson’s trichrome for identification of myocardium, fibrosis, and interstitial space. Samples were examined and imaged using an EVOS XL light microscope (Thermo Fisher Scientific). A custom MATLAB code was used to quantify the percent volume of myocytes, collagen, and interstitial space in the images. To determine statistical differences a Kruskal-Wallis one-way analysis of variance was performed between the control group (n = 3) and the animals implanted with devices (n = 3).

## Acknowledgements

Z. C., N. B., Z. L., and R. T. Y. contributed equally to this work. We thank The George Washington University Nanofabrication and Imaging Center for its facilities regarding device fabrication. L. L. and I. R. E. acknowledge the support of the National Science Foundation (ECCS 2011093) and National Institutes of Health (R21HL152324). L. L. acknowledges support from The George Washington University CDRF and UFF funds. I. R. E. acknowledges Leducq Foundation grant RHYTHM and National Institutes of Health grants (3OT2OD023848 and R01HL141470). R. T. Y. was supported by the American Heart Association Predoctoral Fellowship (19PRE34380781).

