## Supplemental Information for "Flexible and transparent silver nanowire structures for multifunctional electrical and optical biointerfacing"

Z. Chen, N. Boyajian, Z. Lin, R. T. Yin, S. N. Obaid, J. Tian, Dr. J. A. Brennan, Dr. S. W. Chen, A. N. Miniovich, L. Lin, Prof. I. R. Efimov, Prof. L. Lu
Department of Biomedical Engineering
The George Washington University, Washington, DC 20052, USA

Y. Qi, Prof. X. Liu
Department of Civil and Environmental Engineering
The George Washington University, Washington, DC 20052, USA

†These authors contributed equally.

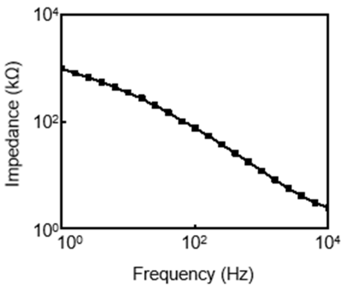

**Figure S1**. Impedance plot of opaque solid Ag films-based microelectrodes.

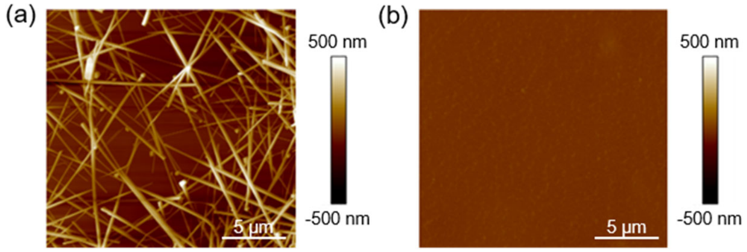

**Figure S2**. AFM images of Ag NWs (a) and solid Ag (b) films.

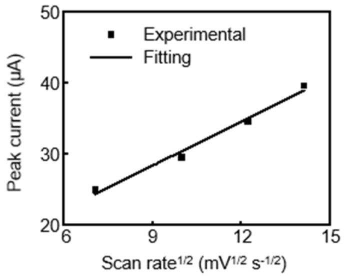


**Figure S3**. Peak current versus square root of scan rate.

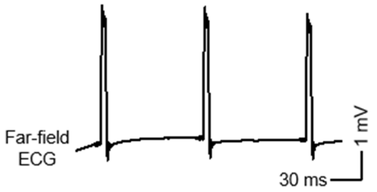


**Figure S4**. Far-field ECG results from the reference electrode during optogenetic pacing at 10 Hz.
